# Transcription Represses Origin Activity in a Late-Replicating Fragile Site

**DOI:** 10.1101/2025.08.01.668051

**Authors:** Juliette Mandelbrojt, Caroline Tonnerre-Doncarli, Aurélie Masson, Théo Baret, Michelle Debatisse, Marie-Noëlle Prioleau

## Abstract

Genome-wide analyses in vertebrates suggest that transcription can negatively regulate replication origin activation, thereby shaping the global landscape of replication initiation. In this study, we investigated how transcription and late replication influence replication origin activity by inserting an efficient minimal model origin into the avian DMD common fragile site (CFS) and a second transcriptionally silent late-replicating region. The DMD gene, which is neither transcribed nor fragile in wild-type DT40 cells, became fragile following transcriptional activation in a genetically engineered DT40 cell line. Previous molecular combing experiments have shown that transcription represses origin firing within the 500 kb 5’ half of the gene. Here, we demonstrate that when a minimal origin is inserted into this repressive region, it remains partially active under basal transcription conditions but is fully inactivated upon transcriptional induction. In contrast, inserting the efficient β-actin promoter/origin at the same site retains origin functionality despite transcriptional activation. Introducing these two origins into a late-replicating, transcriptionally silent locus reveals the same functional hierarchy. Notably, this contrasts with their comparable activity in a mid-late replicating region. These results define a class of replication origins capable of maintaining functionality across diverse chromosomal contexts. Our findings support the model in which CFSs arise due to transcription-dependent repression of replication origin initiation across large, late-replicating genes. Moreover, our results reinforce the idea that replication initiation in late-replicating genomic compartments is intrinsically less efficient than earlier-replicating regions.

## Introduction

The machinery responsible for replicating the vertebrate genome operates on a template organised into distinct compartments, including early- and late-replicating domains, each posing distinctive challenges^1,2^. During the licensing period, from late M to the end of G1, two replicative DNA helicases, Mcm2-7, are assembled in their inactive form on chromatin to form the pre-replicative complex (pre-RC)^3,4^. In S phase, a highly dynamic process involving a cascade of protein recruitment orchestrated by specific regulatory factors, leads to the formation of two CMG helicases (Cdc45.Mcm2-7.GINS)^5,6^. Their activation leads to DNA unwinding and the initiation of bidirectional replication. Thus, the spatiotemporal pattern of DNA replication results from multiple layers of constraints operating throughout G1 and S phase. The late-replicating compartment is mainly characterised by closed and inactive chromatin, posing significant barriers to both pre-RCs deposition and subsequent activation. Specialised trans-acting factors, like the ORC-associated protein (ORCA), are involved in the facilitation of origin licensing in this closed compartment^7–9^. A well-studied exception to gene paucity of late-replicating domains is the presence of a series of large genes (>300 kb) hosting common fragile sites (CFSs)^10,11^. CFSs are chromosomal regions that remain incompletely replicated when cells submitted to replication stress enter mitosis^10^. Importantly, CFSs have been shown to be tissue-specific hotspots of chromosome rearrangements associated with cancer progression^12^. Several hypotheses have been proposed to explain how transcription delays the completion of replication at CFSs. One postulates that R-loop formation upon head-on encounter of the replication and the transcription machines blocks replication fork progression and induces fork collapse^13,14^. However, this model is challenged by evidence showing that, in gene-dense early replicating regions, initiation events occur within actively transcribed genes, mostly in their 5’ part^15,16^. In addition, copy number variants (CNVs) found in cancer cells cluster within CFSs, but not within regions enriched in highly transcribed genes^14^. Furthermore, depletion of RNase H1, a major contributor to R-loop resolution, does not impact CNVs frequency or distribution. Finally, DRIP-seq analyses showed that CFSs are not enriched in R-loops. Together, these results suggest that R-loops are not the primary driver of transcription-induced CFS instability^17^. A second model relies on the observation that the core of two canonical CFSs is depleted in replication initiation events. This suggests that ongoing transcription actively removes or displaces pre-RCs^18,19^, leaving these regions more prone to incomplete replication when replication forks slow down. In addition, genome-wide analyses further show that transcription units tend to be depleted in replication initiation events, regardless of replication timing^20,21^. This observation suggests a general mechanism by which ongoing transcription would remove the pre-RCs, or prevent their setting and further steps of origin building in consequence. The second hypothesis was supported by *in vitro* and *in vivo* studies in S. *cerevisiae*, showing that the transcriptional machinery could indeed displace pre-RCs, though this displacement was limited to a few kilobases^22^. *In vivo* experiments, where the well-studied S. *cerevisiae* ARS1 element was artificially inserted into a transcription unit, showed inactivation of the origin^23^.

Despite these insights, how transcription modulates the behaviour of efficient vertebrate replication origins across different chromosomal contexts remains poorly explored. However, although sharing many similarities, the nature of the replication machinery (*trans*-factors) and of the replication initiation sites (*cis*-elements) in yeast and vertebrates displays specificities that may lead to different behaviours *in vivo*. Moreover, vertebrate genomes feature more complex chromatin landscapes and regulatory layers, underscoring the need for dedicated *in vivo* studies. In this study, we utilised a well-characterised chicken minimal origin of replication devoid of intrinsic transcriptional activity to investigate how this *cis*-element functions in two late-replicating chromosomal contexts, one embedded within a transcription unit and the other in a silent context. We further compared its activity to that of an efficient origin located within the constitutive β-actin promoter. Our results demonstrate that transcriptionally active late-replicating chromatin represses the activity of the minimal origin, with repression levels correlating with transcriptional output. In contrast, the β-actin promoter/origin remains efficient in all tested conditions. This study allows dissection of the molecular interplay between transcription, replication origin activation, and chromatin environment, advancing our understanding of fragile site biology.

## Results

### Ongoing transcription across the late-replicating DMD gene suppresses firing of the β-globin minimal origin

To determine whether ongoing transcription within large genes can suppress replication origin firing in DT40 cells, we analysed the behaviour of the β^A^-globin minimal origin inserted into appropriately engineered DMD gene. Indeed, we took advantage of a cell line in which both alleles of this late-replicating and 1 Mb long gene, which is naturally silent and non-fragile in wild type (WT) DT40 cells, have been replaced by the tetracycline-inducible promoter (DMD^Tet/Tet^)^19^. In the absence of tetracycline treatment, these cells exhibit low levels of transcription and gene fragility, two features that were accentuated upon tetracycline treatment. A previous molecular combing analysis has revealed that in WT cells, initiation sites are randomly distributed across the gene. However, in tetracycline-treated DMD^Tet/Tet^ cells, initiation sites are excluded from a region extending over approximately 500 kb within the 5’ part of the gene resulting in a large termination zone (Figure 1A). Despite transcriptional activation, the core of the gene remained late replicating and the termination zone was shown to host the fragile region^19^. Given that replication initiation exclusion is limited to the first half of the DMD gene, we wondered if the transcription rate was similar in the 5’ and 3’ regions of this long gene. To address this, we labelled asynchronous DMD^Tet/Tet^ cells with EU for 15 minutes. After the capture of newly synthesised RNA and reverse transcription, the nascent transcript levels were analysed by EU-seq. Notably, we observed a marked decrease in the number of nascent transcripts in the second half of the gene, which corresponds to the region where initiation events are preserved (Figure 1B). These findings support the model positing that transcription-induced remodelling of replication origin distribution prevents replication completion of large genes before mitosis upon fork slowing, resulting in fragility. To directly assess the impact of passage of the transcription machinery on origin activity in a controlled setting, we employed a previously characterised efficient β^A^-globin minimal origin devoid of promoter activity^24^. This 90 bp minimal origin, which contains two potential G-quadruplexes motifs on the same DNA strand (dimeric pG4, Figure 1B), was inserted by homologous recombination, into the centre of the DMD termination zone on one chromosome in DMD^Tet/Tet^ cells (Figure 1B). Given that approximately 30% of efficient origins in human and chicken cells share this organization, the minimal origin serves as a representative model for efficient replication initiation sites. To prevent premature transcription termination caused by the SV40 poly(A) signal in the construct, the active minimal origin was inserted with G-tracks aligned to the template strand (Fig. 1B). Quantification of the small nascent strand (SNS) relative enrichment revealed that in the absence of tetracycline induction, the minimal origin remains functional, albeit its efficiency decreases significantly (ninefold) when compared with previously tested insertion in mid-late replicating region (Figure 1C). Supporting a model in which transcription suppresses initiation events, we observed an additional sixfold decrease in initiation efficiency upon tetracycline addition (Figure 1C). To confirm the specificity of the SNS relative enrichment signal detected in the absence of tetracycline induction, we analysed a control cell line containing an inactive version of the minimal origin at the same locus (Figure 1B)^24^. In this control, SNS relative enrichment was tenfold lower than with the active minimal origin (Figure 1C). Overall, these results strongly support the hypothesis that transcription inhibits the initiation at an efficient origin.

**Figure 1.**
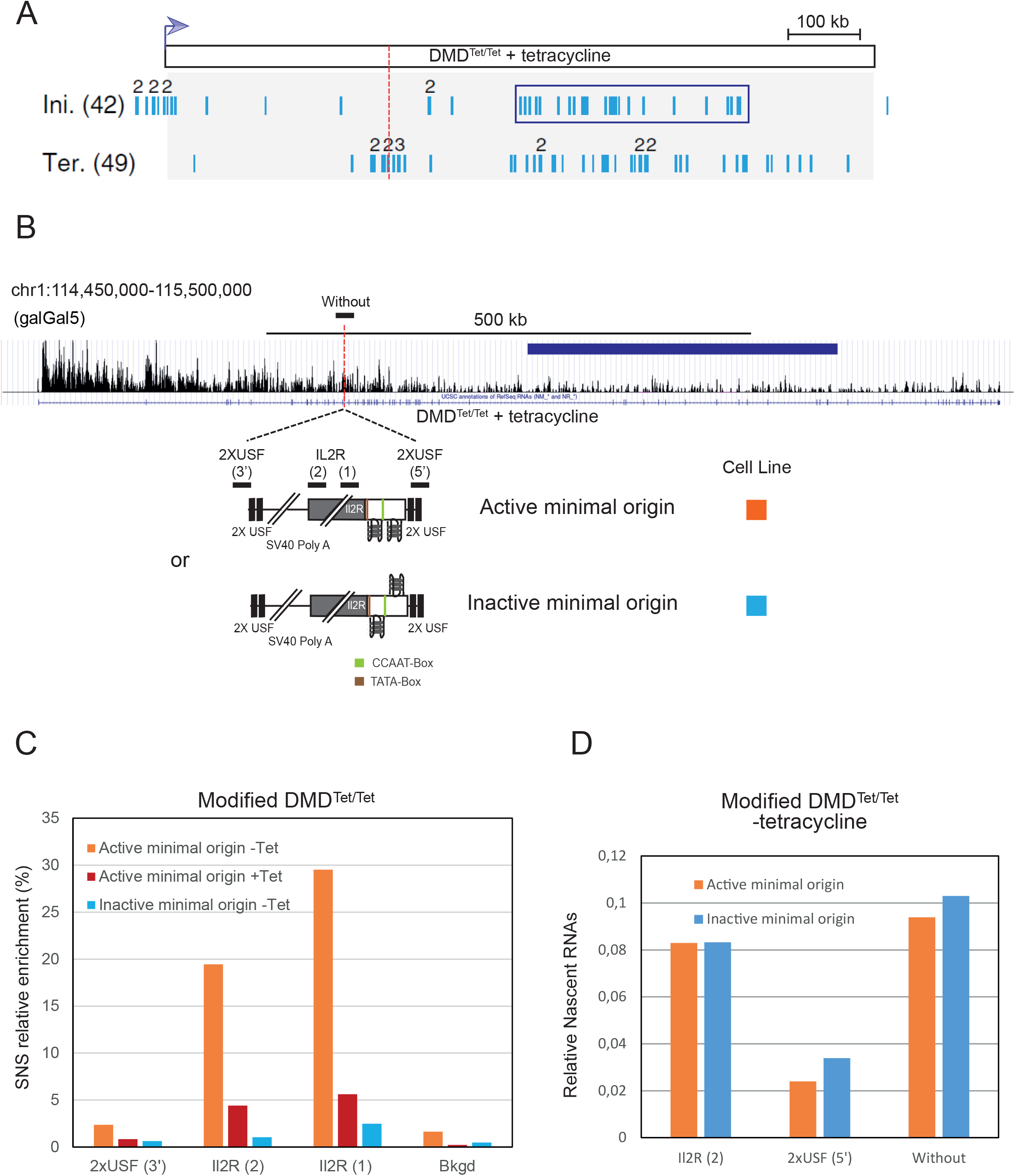
Transcription through the DMD^Tet/Tet^ fragile site represses origin firing. (A) Schematic representation of replication initiation (Ini) and termination (Ter) events identified by molecular combing in DMD^Tet/Tet^ cells grown in presence of tetracycline (Blin *et al*, 20219). The insertion site for the active or inactive minimal origin is indicated by a red dotted line. The blue rectangle surrounds a zone of initiation, which is also represented in panel B. (B) UCSC genome browser view showing EU-seq signal across the DMD gene in the DMD^Tet/Tet^ cells grown in the presence of tetracycline. Although the studied cell line is devoid of insertion, the insertion site for the active or inactive minimal origin is indicated by a red dotted line. Below schematic representations of the inserted constructs are shown: each construct is flanked by two USF (Upstream Stimulating Factor) binding sites and contains a 90 bp minimal origin composed of two pG4s on the same strand (dimeric pG4). The second one is surrounded by a CCAAT and a TATA box represented in green and brown respectively. In the active origin, the G-tracks are on the template strand. The inactive minimal origin is similar to the active one, except that the first pG4 is on the reverse complement strand. Black lines above the construct indicate the positions of amplicons used for SNS and nascent RNA quantifications, specific to the modified chromosome. The amplicon above the DMD gene overlaps the insertion site and is specific to the unmodified chromosome. (C) SNS relative enrichments along the ectopic active minimal origin obtained in clones (n=2) cultured without (orange bars) and with (red bars) tetracycline or along the inactive minimal origin in clones (n=2) cultured without tetracycline (blue bars). The amplicons used for quantification are indicated in panel B. Bkgd refers to an amplicon located 5 kb away from the insertion site. An amplicon within the endogenous r-origin was arbitrarily set at 100% to calculate the SNS relative enrichment. (D) Relative abundance of nascent RNAs at and around the minimal origin. Nascent transcripts were measured in two cell lines (n=2) containing the active (orange bars) or the inactive (blue bars) minimal origin within the DMD^Tet/Tet^ cell line, grown without tetracycline. Nascent transcripts levels at the endogenous Bu1a gene were arbitrarily set at 1 for normalization. The amplicons used for RT-qPCR quantification are shown in panel B.

### Transcription progresses unimpeded through an active origin in a late replicating region

To determine whether the presence of an efficient origin affects RNA polymerase II progression, we quantified the amount of nascent transcripts immediately downstream of the minimal origin (Figure 1D). We used an amplicon overlapping the insertion site to quantify nascent transcripts on the unmodified chromosome (amplicon Without, Figure 1B). We compared this expression with that observed on the modified chromosome both upstream and downstream of the minimal origin (IL2R (2) and 2XUSF (5’) respectively, Figure 1B). As an additional control, we also analysed two cell lines in which the inactive minimal origin was inserted at the same site. As a reference, we quantified the transcripts from the Bu1a gene, arbitrarily setting its expression level to one. To ensure sufficient cDNA yields for qPCR analysis, nascent RNAs were labelled with EU for one hour as previously described^19^. Thus, the measured RNA levels reflect both the transcriptional activity and the stability of the analysed transcripts. To minimize the impact of the construct insertion on RNA synthesis, we introduced the construct within an intron. We observed a two-fold decrease in nascent transcript levels downstream of the minimal origin (Figure 1D). This reduction was consistent for both active and inactive origins, suggesting that the presence of pG4 structures may impede RNA polymerase II progression in this specific chromosomal context. RT-qPCR analysis of nascent RNAs revealed that, although the signal after the origin was reduced, it remained clearly above background, as no amplification was observed in the negative control lacking reverse transcriptase (Supplementary Figure 1A). This suggests that RNA polymerase progression through the origin occurs. To compare the transcriptional activity of the DMD gene in its uninduced and induced states, we analysed nascent transcript levels, labelled for 15 minutes with EU, across the entire gene by EU-seq. We observed an approximately 2.5-fold increase in transcription between the -Tet and +Tet conditions, consistent with our RT-qPCR results and previously published data (Supplementary Figure 1B)^19^. Overall, our results support the model proposing that transcriptional progression through the long DMD gene inhibits origin firing while firing of an intragenic origin does not affect transcription of this large and late-replicating gene.

### The ‘-actin origin/promoter remains efficient when inserted into the DMD^Tet/Tet^ gene

To further investigate how transcription affects origin firing within the DMD gene, we analysed cell lines containing the minimal inactive origin before excision of the strong β-actin promoter/origin, which drives the selection gene expression (Figure 2A). This β-actin promoter contains nine pG4s (six of which are within dimeric pG4s). Consistent with the model proposing that dimeric pG4s are efficient replication initiation sites, SNS-seq analysis in WT DT40 cells revealed a pronounced enrichment zone covering the endogenous β-actin promoter (Figure 2B). Previous work demonstrated that a 1264 bp fragment encompassing this promoter induces efficient replication initiation when ectopically inserted in a mid-late replicating region^25^. Accurately assessing the global activity of this ectopic origin by SNS relative enrichment is challenging due to the presence of the endogenous β-actin locus. We therefore assessed the impact of this origin on replication timing (RT), as a shift in RT can reflect origin efficiency, as previously observed^24–27^. RT shift analysis of two independent cell lines indicated that the ‘-actin origin alone is sufficient to induce a shift toward earlier replication when cells were grown without tetracycline, although the region remains replicated in the second half of S phase (Figure 2C). This contrasts with the incapacity of the minimal origin to shift the RT under similar conditions but aligns with the low SNS relative enrichment observed at the minimal origin (Supplementary Figure 1C and Figure 1C). To further explore the capacity of the β-actin promoter/origin to remain efficient when transcription is up-regulated, we also test its capacity to advance RT upon tetracycline induction. We observed no significant difference in RT shift with or without tetracycline, suggesting that the ‘-actin origin maintains its firing capacity despite transcription induction (Figure 2D). These results show that the efficiency of the origins of replication depends largely on the chromosomal context in which they are found. While the minimal origin and the ‘-actin origin exhibit similar efficiencies, based on RT shifts, in a region replicated in the mid-late S phase^25^, the ‘-actin origin is more efficient within a late replication region and is also able to remain functional when crossed by RNA polymerase II. On the other hand, the activity of the minimal origin is greatly affected in a late replicating context, with an increase in the transcription rate leading to its complete inhibition.

**Figure 2.**
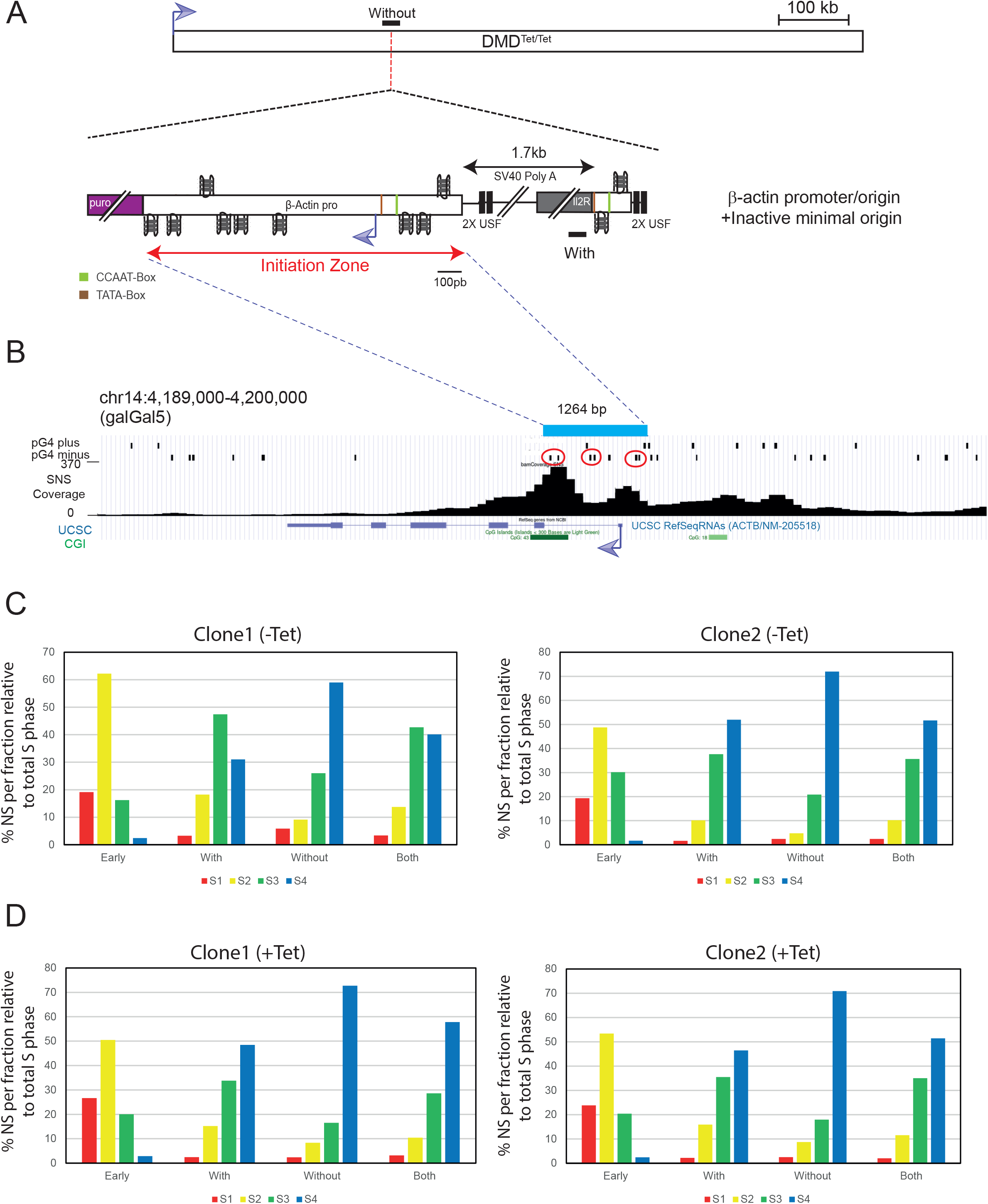
The ‘-actin promoter/origin is not repressed when inserted within the transcribed DMD gene. (A) Schematic representation of the DMD^Tet/Tet^ cell line containing the ‘-actin promoter/origin fused to the minimal inactive origin. Black lines indicate the position of amplicons used for allele specific quantifications of RT profiles. The red dotted line indicates the site of insertion. (B) The endogenous ‘-actin promoter is located within a replication initiation zone. The top blue bar defines the 1264 bp region derived from the ‘-actin promoter present in the construct. Below pG4 detected on the plus and minus strand in Zheng *et al*, 2020^33^ are shown. Dimeric pG4s found within the ectopic ‘-actin promoter are surrounded by red ellipses. CpG islands (CGI) and UCSC RefSeq RNAs annotations are also indicated at the bottom in green and blue respectively. (C) Replication timing (RT) profiles of each chromosomal allele were determined after targeted transgene integration, using the allele-specific analysis by quantitative PCR. BrdU pulse-labelled cells were sorted into four S-phase fractions, from early to late (S1 to S4) and the immuno-precipitated newly synthesized strands (NS) were quantified by qPCR in each fraction. Specific primer pairs determine the RT profile for the modified allele (With), the WT allele (Without) and 5 kb away from the insertion site on both alleles (Both). The endogenous β-globin locus was analysed as an early-replicated control (Early). Two independent clones containing the indicated constructs were analysed. (D) Same experimental procedure as in C, except that the cells were grown for 24 hours in the presence of tetracycline before RT analysis.

### Replication origin efficiency hierarchy is conserved in a silent, late-replicating domain

We tested the effect of transcription passing through a minimal efficient origin and the ‘-actin promoter/origin in a late-replicating common fragile site. We observed different behaviours, demonstrating that the chromosomal context imposes specific constraints on the replication machinery. In order to test the impact of a late replication context without transcription, we initially tried to insert the minimal origin in the wild-type DMD context. Despite numerous attempts, we were unable to obtain clones that had inserted the construct by homologous recombination, likely due to the poor chromatin accessibility in this silent region which impairs efficient homologous recombination. As an alternative, we tested a late-replicating region for which we had already produced several clones, showing that homologous recombination is feasible in this silent and late-replicating region. The timing of replication is comparable to that found in the DMD gene, and the locus was shown to be tightly bound to the nuclear periphery and to have a tight late replicating control^28^. In this late replicating region, SNS relative enrichment analysis showed that the minimal origin was repressed compared to the mid-late replicating domain, but exhibited a level of SNS relative enrichment similar to that observed when inserted in the DMD gene of the DMD^Tet/Tet^ cell line and in absence of tetracycline induction (Figure 3B, graph on the left compared with Figure 1C, orange bars). This result aligns with the absence of any replication timing shift toward earlier replication of the modified chromosome (Figure 3C). As a control, we quantified SNS relative enrichment in a cell line containing the inactive minimal origin adjacent to the active ‘-actin promoter/origin. SNS relative enrichment at the inactive minimal origin was approximately fivefold lower than at the active minimal origin (Figure 3B, on the right, enrichment at IL2R (1 and 2) is fivefold lower) confirming the weak but detectable activity of the active minimal origin at this locus. However, we observed a more pronounced SNS relative enrichment at the ectopic ‘-actin origin. As previously mentioned, accurately assessing the global activity of this ectopic origin is challenging due to the presence of the endogenous copy. RT shift analysis of this cell line indicated that the ‘-actin origin is sufficient to induce a shift toward earlier replication, although the region remains late-replicating (Figure 3D). The extent of this shift is comparable to that seen in the transcribed DMD gene, suggesting that the β-actin origin is active in a substantial fraction of cells and functions as an efficient origin in this repressive context. Overall, this chromosomal context behaves similarly to the uninduced DMD^Tet/Tet^ gene, exerting a global repressive effect on origin activity of the minimal origin, although the β-actin promoter/origin retains its capacity to advance RT. This result underscores the significant influence of the chromosomal environment on origin function. It also suggests that a strong promoter, associated with efficient replication origins containing multiple dimeric pG4 motifs, establishes a chromatin context around the promoter that is highly conducive to origin activation.

**Figure 3.**
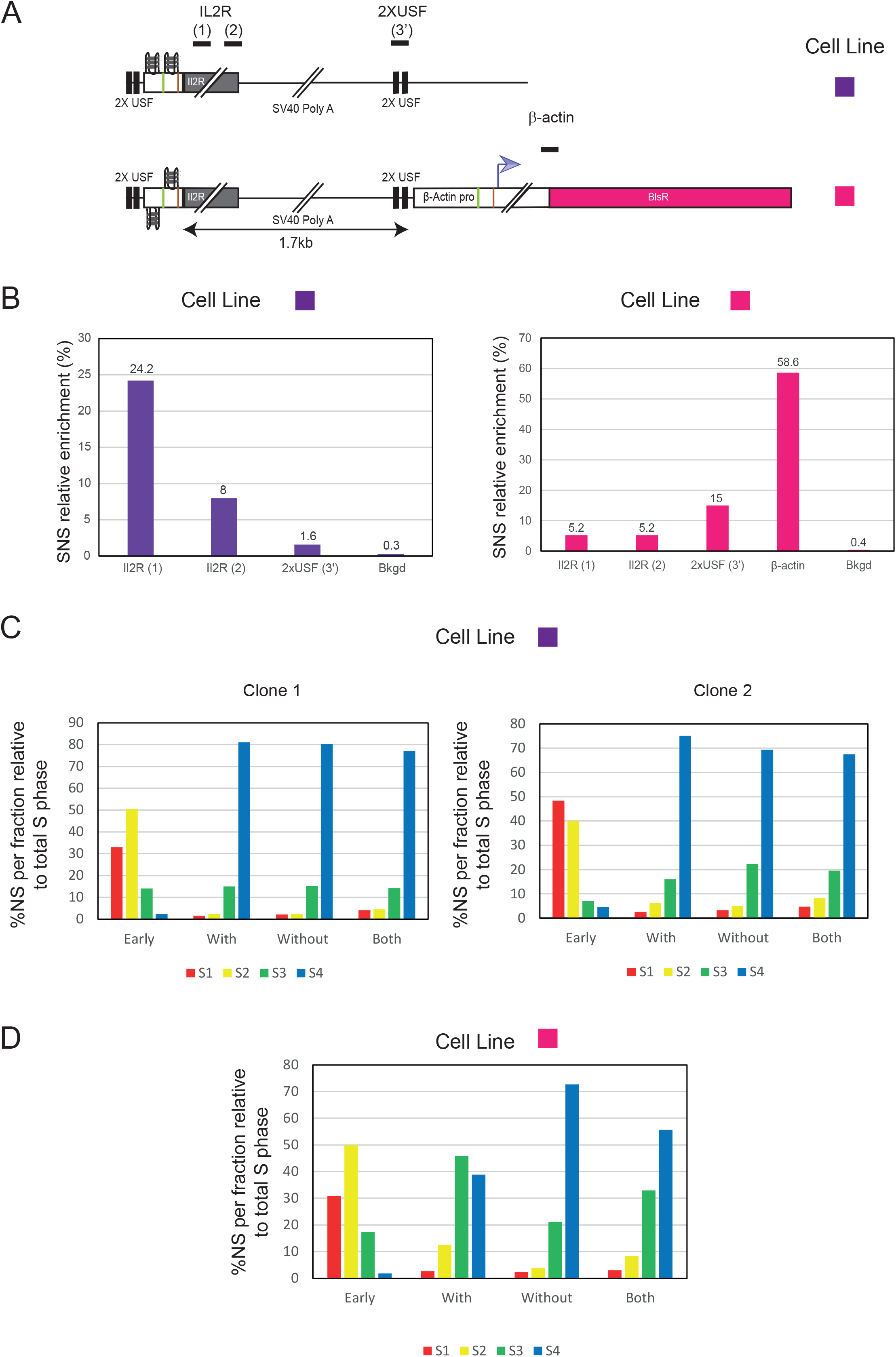
A late chromosomal environment represses the minimal efficient origin but not the ‘-actin promoter/origin. (A) Schematic representation of the two analysed cell lines. Both constructs were inserted at the Late2 locus (chr1:177,936,192 bp, galGal5) previously described in Duriez *et al*, 2019 and Brossas *et al*, 2020. One cell line harbours the active minimal origin alone (purple), while the other contains the inactive minimal origin fused to the ‘-actin promoter/origin (pink). Black lines above the construct indicate the position of amplicons used for SNS quantifications. (B) SNS relative enrichments along the ectopic active or inactive minimal origin and the ‘-actin promoter/origin obtained in clones (n=2), the mean values are indicated above each bar. The amplicons used for quantification are indicated below each graph. Bkgd refers to the amplicon located 11 kb away from the integration site. One amplicon within the endogenous r-origin was arbitrarily set at 100% to quantify the relative SNS abundance. (C) Replication timing (RT) profiles of each chromosomal allele were determined after targeted transgene integration, using the allele-specific analysis by quantitative PCR. Specific primer pairs determine the RT profile for the modified allele (With, amplicon IL2R (1)), the WT allele (Without, amplicon overlapping the site of insertion) and 3.5 kb away from the insertion site on both alleles (Both). The endogenous β-globin locus was analysed as an early-replicated control (Early). Analyses of two independent clones containing the active minimal origin inserted at the Late2 locus. (D) RT profiles for one clone containing the minimal inactive origin next to the ‘-actin promoter inserted at the Late2 locus are shown.

## Discussion

Transcription and replication machineries share the same DNA template, making their coordination inherently complex. This complexity is heightened by the temporal separation between the assembly of pre-replication complexes (pre-RCs) and the initiation of replication. During S phase, pre-RCs displaced by transcription can be reloaded dynamically^29^. However, once S phase begins, new pre-RCs can no longer be assembled. Therefore, their displacement by transcription leads to a blockade of replication initiation within actively transcribed regions. Pre-RCs activated at the onset of S phase face only a short window during which transcriptional interference can occur, whereas those activated later in S phase are exposed to a much longer period of potential disruption. To investigate how transcription affects replication origins in late-replicating contexts *in vivo*, we compared a minimal 90 bp efficient origin devoid of promoter activity to a longer 1264 bp origin containing a functional promoter. Our results show that the chromosomal environment of a long, transcriptionally active, late-replicating gene corresponding to a common fragile site (CFS), strongly represses the activity of the minimal efficient replication origin (Figure 4). This observation aligns with previous findings from molecular combing studies, which have revealed a depletion of replication initiation events in the cores of CFSs, such as FRA3B in human cells and DMD in chicken cells, in a transcription-dependent manner. However, direct experimental evidence that transcription can inactivate an efficient replication origin has been lacking. A prior study in avian DT40 cells reported a reduction of initiation events in the first half of the transcribed DMD gene. Here, our work demonstrates that this depletion correlates with nascent transcription density, suggesting a direct relationship between transcriptional activity and repression of local origin firing. To further probe this relationship, we assessed the activity of an efficient origin inserted into the transcriptionally depleted region under conditions that induce gene fragility. In the absence of tetracycline-induced transcription, the minimal origin remained functional, albeit repressed. However, upon induction, transcription inactivated the origin. Notably, the repression of origin activity increased in tandem with elevated transcription and fragility of the DMD gene, pointing to a causal relationship. A previous study using the same inducible cell line showed that tetracycline treatment increased the proportion of chromosome breaks from 13% to 48% (Figure 4)^19^. Our finding that origin inactivation accompanies this increase in fragility supports the notion that CFS instability results from the transcription-dependent suppression of replication initiation events by RNA polymerase II (Figure 4). Interestingly, the strong β-actin promoter/origin remained efficient even under induced DMD transcription. This finding highlights a functional hierarchy among replication origins, with some origins capable of functioning independently of their chromatin or transcriptional environment. We observed that the β-actin origin contains multiple G-quadruplexes (pG4s), including three pG4 dimers, which may be essential for the formation of a robust origin. Origins of this type are likely well represented among the constitutive or core origins described in Picard et al^30^ and Akerman et al^15^, respectively. Previous work has shown that such pG4 dimers are critical for the activity of the minimal origin and are found in approximately 30% of efficient origins in both human and chicken cells^24^. However, we did not genetically dissect the 1264 bp β-actin origin to define which *cis*-elements are required for its function. Our findings suggest that such elements are depleted from CFSs, consistent with the AT-rich nature of these genomic regions. We also examined the impact of a late-replicating chromatin context lacking transcriptional activity on the function of both the minimal and β-actin origins. Previously, we demonstrated that the minimal origin is efficient within a mid-late S phase region, as evidenced by strong relative enrichment of short nascent strands (SNSs) and a significant advance in local replication timing^24^ (Figure 4). In this context, the β-actin origin exhibited similar behaviour. However, in a silent, late-replicating region, the minimal origin shows modest SNS enrichment and induces no significant change in replication timing reflecting reduced origin activity. In contrast, insertion of the β-actin origin into the same region resulted in a pronounced advance in replication timing mirroring our results in the transcribed DMD gene. This supports the existence of a functional hierarchy among efficient origins, likely defined by the presence and arrangement of dimeric pG4 *cis*-elements. Moreover, this hierarchy appears to be context-dependent: in a more permissive (mid-late) region, both types of origins perform similarly, whereas in a more restrictive (late-replicating) context, the minimal origin is repressed while the promoter-associated β-actin origin remains active. These results underline the critical role of *cis*-elements in the establishment of the spatiotemporal replication programme, while also highlighting the role of RNA polymerase II passage in the establishment of regions depleted in replication initiation events. In this study, we focused on a late-replicating, transcriptionally active region. Further investigations are required to assess the impact of the high transcriptional activity observed in early-replicating regions on replication origin firing. In addition to common fragile sites, early-replicating fragile sites (ERFSs) constitute a distinct class of fragile genomic regions localised in highly transcribed, gene- and R-loops rich domains that replicate early in S phase^31^. This organisation may promote genotoxic head-on (HO) collisions between the DNA replication machinery and RNA polymerase^32^. Transcription-replication interference emerges as a fundamental mechanism driving fragile sites formation throughout S phase, although the mechanisms involved in the early and late S-phase are quite different. Therefore, exploring the behaviour of our minimal origin model in an early replicating region would provide a more comprehensive understanding of how transcriptional activity, replication timing, and the chromatin environment collectively influence origin function and genome stability.

**Figure 4:**
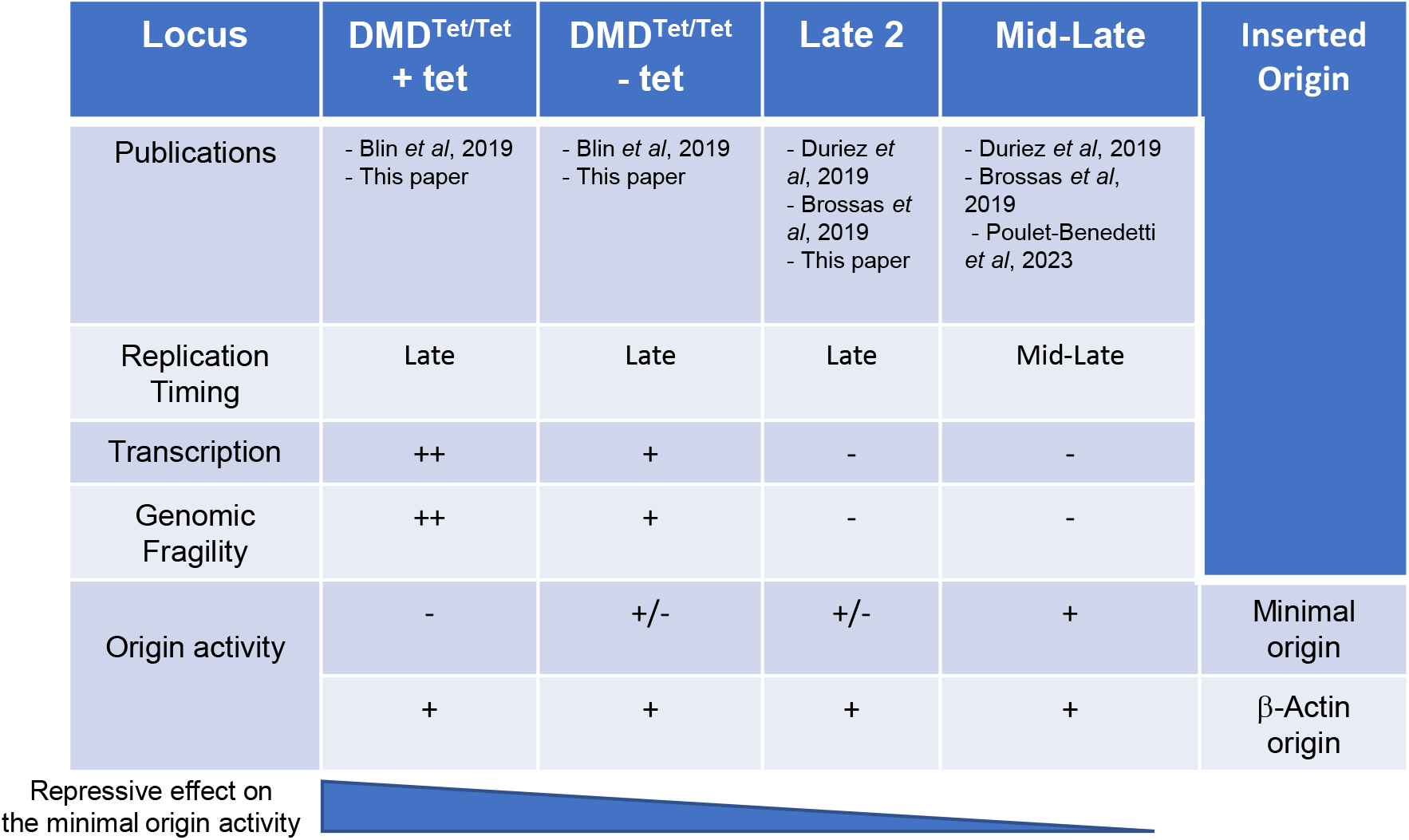
A summary of the results obtained from inserting the minimal and β-actin origins at three different loci. The publications describing the results relevant to this study are indicated in the top row. In the DMD^Tet/Tet^ cell line, induction with tetracycline leads to a two-to threefold increase in nascent RNAs and a fourfold increase in the % of broken chromosomes in metaphase. Origin activity was assessed using two assays: (1) the short nascent strand (SNS) relative enrichment and (2) the origin’s ability to locally advance the replication timing (RT). A ‘+’ indicates a high level of SNS relative enrichment and local RT advancement. A ‘+/-’ indicates a low SNS relative enrichment without local RT advancement. A ‘-’ indicates a SNS relative enrichment comparable to background and no detectable RT advancement.

## Methods

### Plasmid construction

The targeting vectors used for homologous recombination in DT40 cells were constructed with the multisite Gateway Pro kit (Thermo Fischer Scientific #12537100). Vectors containing the 5’ and 3’ target arms used for specific recombination at the Late 2 (chr1:177,936,192 bp, galGal5) were described previously^28^. New arms were prepared for specific targeting in the DMD^Tet/Tet^ gene (chr1:114.796.392-114.798.322, galGal5). The 5’ and 3’ target arms for homologous recombination were amplified from DT40 genomic DNA with primer pairs listed in Supplementary Figure 4. The arms were oriented so that the minimal origin construct was inserted with the pG4s on the template strand for the active origin version. We used four entry vectors to generate the new minimal origin construct inserted at either the DMD or Late 2 sites: two entry vectors containing the 5′ and 3′ target arms for specific insertion, one entry vector pDONR221 P5-P4 containing the minimal origin described previously^24^ and one entry vector pDONR221 P4r-P3r containing the drug selection cassette (β-actin-PuroR at the DMD locus or β-actin-BsR at the Late 2 locus) described previously^26^. Final vectors were generated by recombining compatible *att* sites between the entry vectors, with LR clonase (Thermo Fisher Scientific #12538120). For electroporation, the final vectors were linearized with *ScaI* (NEB #R3122S).

### Cell culture condition and transfection

DT40 cells were grown in RPMI 1640 medium supplemented with Glutamax (Thermo Fisher Scientific #61870010), containing 10% FBS, 1% chicken serum, 0.1mM β-mercaptoethanol, 200U/mL penicillin, 200μg/mL streptomycin and 1.75 μg/mL of amphotericin B at 37°C, under an atmosphere containing 5% CO_2_. Cells were electroporated as previously described. Cell clones were selected on media containing a final concentration of 20 μg/ml blasticidin or 1μg/mL puromycin. Genomic DNA was extracted from cells in lysis buffer (10 mM Tris pH8.0; 25 mM NaCl; 1 mM EDTA and 200 μg/mL proteinase K). Clones into which the plasmid DNA was integrated were screened by PCR with primer pairs designed to bind on one side of the insertion site such that one primer bound within the construct and the other primer bound just upstream or downstream from the arm used for recombination (Supplementary Fig. 2 and Supplementary Figure 4). The *BsR* or PuroR resistance gene was excised from positive clones using the Cre-LoxP system. DT40 cells constitutively express a tightly regulated Cre recombinase fused to a mutated oestrogen receptor (Mer). This inactive Mer-Cre-Mer fusion protein can be transiently activated in the presence of 4-hydroxytamoxifen, resulting in the efficient excision of genomic regions flanked by two recombination signals (*loxP* sites) inserted in the same direction. For the excision of the genomic DNA flanked by loxP sites, we treated 3 × 10^5^ cells with 5 μM 4-hydroxytamoxifen (Sigma Aldrich #T176) for 24 h. Subclones were obtained by plating dilutions of the treated cell suspension at a density of 50, 150, and 1500 viable cells per 10 ml in 96-well flat-bottomed microtiter plates. We assessed the excision of the β-actin-BsR or β-actin-PuroR selection cassette by growing cells in the selective media for 72 hrs, with a control plate without selection. Cell clones that died in the presence of the selection drug were selected for the correct excision. Transcriptional activation of the DMD^Tet/Tet^ gene was achieved by treating cells with 1μg/mL tetracycline for 24hrs.

### SNS purification

SNS purification was performed as previously described with some slight modifications. Fresh or frozen cultured cells were used for total genomic DNA extraction and the T4 polynucleotide kinase (Biolabs #M0201S) concentration was adjusted to 100U and incubated for 30 min at 37°C. Proteinase K (Thermo Fischer Scientific #EO0491) digestion was realized at a final concentration of 625μg/ml for 30min at 50°C.

### Replication Timing analysis

For RT experiments, about 10^7^ exponentially growing cells were pulse-labelled with 5-Bromo-2’-deoxyuridine (BrdU, Sigma-Aldrich #B9285) for 1 h and sorted into four S-phase fractions, from early to late S phase. The collected cells were treated with lysis buffer (50 mM Tris pH 8.0; 10 mM EDTA pH 8.0; 300 mM NaCl; 0.5% SDS, 0.2 mg/ml of freshly added proteinase and 0.5 mg/ml of freshly added RNase A), incubated at 56°C for 2 h and stored at -20°C, in the dark. Genomic DNA was isolated from each sample by phenol-chloroform extraction and alcohol precipitation and sonicated four times for 30s each, at 30s intervals, in the high mode at 4°C in a Bioruptor water bath sonicator (Diagenode), to obtain fragments of 500 to 1000 bp in size. The sonicated DNA was denatured by incubation at 95°C for 5 minutes. We added monoclonal anti-BrdU antibody (BD Biosciences #347580) at a final concentration of 3.6 μg/ml in 1x IP buffer (10 mM Tris pH 8.0, 1 mM EDTA pH 8.0, 150 mM NaCl, 0.5% Triton X-100, and 7 mM NaOH). We used 50 μl of protein-G-coated magnetic beads (from Thermo Fisher Scientific #10004D) per sample to pull down the anti-BrdU antibody. Beads and BrdU-labelled nascent DNA were incubated for 2-3 hours at 4°C, on a rotating wheel. The beads were then washed once with 1x IP buffer, twice with wash buffer (20 mM Tris pH 8.0, 2 mM EDTA pH 8.0, 250 mM NaCl, 0.25% Triton X-100) and then twice with 1x TE buffer pH 8.0. The DNA was eluted by incubating the beads at 37°C for 2hrs in 250 μl 1x TE buffer pH 8.0, to which we added 1% SDS and 0.5 mg/ml proteinase K. DNA was purified by phenol-chloroform extraction and alcohol precipitation and resuspended in 50 μl TE. The BrdU-labelled nascent strands (NS) quantification was performed as previously described^25–27^.

### Flow cytometry analysis and sorting

After BrdU incorporation, DT40 cells were washed twice with PBS, fixed in 75% ethanol and stored at-20°C. On the day of sorting, fixed cells were resuspended at a final concentration of 2.5 × 10^6^ cells/mL in 2% foetal bovine serum in PBS (Sigma, #CA-630), 50 μg/ml propidium iodide and 0.5 mg/ml RNase A, and incubated for 30 minutes at room temperature. Singlet cells were sorted with a FACSAria Fusion (BD Biosciences). Four fractions of S-phase cells (S1-S4), each containing 5×10^4^ cells, were collected and further treated for locus-specific RT analyses.

### Extraction of 5-ethynyl uridine incorporated nascent RNAs and reverse transcription

Exponentially growing DT40 cells were treated with 0.5mM 5-ethynyl uridine for 15 min or 1 hour. Transcriptional activation of the DMD^Tet/Tet^ gene when required was achieved by treating cells with 1μg/mL tetracycline for 24 hrs prior to EU labelling. 5-ethynyl uridine-labeled RNA was captured using the Click-iT Nascent RNA Capture Kit (Thermo Fischer Scientific, #C10365) according to the manufacturer’s instructions. Total RNA was extracted from 10×10^6^ cells with the miRNeasy kit (Qiagen, #217004). 5 μg of total RNA was biotinylated thanks to the click reaction between 5-ethynyl uridine and azide-modified biotin. 2 μg of biotinylated RNA was then purified using streptavidin magnetic beads. Nascent RNAs were reverse transcribed on the beads (RT+) using Superscript VILO cDNA Synthesis Kit (Thermo Fischer Scientific, #11754050). Negative controls (RT -) were performed with the same procedure but without the addition of reverse transcriptase. The comparison of RT + and RT - samples was used to validate the absence of genomic DNA in the RNA samples. Real-time qPCR was performed using primer pairs listed in Supplementary Table 1. RNA levels were calculated relative to the Bu1a expression level. Nascent RNA quantification experiments were performed using two clonal cell lines per analysed construct.

### Real-time PCR quantification of DNA

Real-time qPCRs were executed according to the MIQE guideline. The LightCycler 480 Real-time PCR system with the SYBR Green I Master Mix (Roche Life Science, # 04887352001) was used for the real-time PCR quantification of BrdU-labeled nascent strands (NS), genomic DNA extracted from clonal cell lines, short nascent strands or 5-ethynyl uridine incorporated nascent RNAs. Each sample was quantified at least in duplicate. For all reactions real-time PCR was performed under the following cycling conditions: initial denaturation at 95°C for 5 minutes, followed by 50 cycles of 95°C for 10 s, 61°C for 20 s, 72°C for 20 s and fluorescence measurement. Following amplification, a thermal melting profile was used for specific amplicon validation.

### Sequencing library preparation

Sequencing libraries were prepared using NEBNext Ultra II Directional RNA Library Prep Kit for Illumina (NEB #E7760S) following the manufacturer’s instructions for RT-nascent RNAs. The samples were not subjected to size selection, but were cleaned up for adaptor-ligated DNA using a SPRISelect Reagent Kit (Beckman coulter #B23317). Libraries were prepared from cDNA produced using 1μg of biotinylated EU-RNA as a starting amount. A 10-fold diluted adaptor was used for the adaptor ligation. Library amplification was performed using the Unique Dual Index UMI Adaptors RNA Set 1 (NEB#E7416) with twelve PCR cycles. The mean size and quality of the library molecules were determined using an Agilent Bioanalyser High Sensitivity DNA chip (Agilent technologies, #5067– 4626).

### Sequencing and data processing

Sequencing was performed by Novogene using an Illumina NovaSeq X Plus sequencer with 10B flow Cell. Samples were sequenced in paired-end mode (150 bp paired-end reads) according to standard procedures. Approximately 10M of reads (∼3Gb) were generated per sample. Paired-end reads were aligned to the Gallus gallus genome assembly galGal5 using bowtie2 (version 2.5.1) with default parameters. Resulting SAM files were processed using Samtools (version 1.21). BAM files were indexed with samtools index. Processed BAM files were converted into BigWig coverage file using the Galaxy France platform for visualisation in the UCSC Genome Browser. Coverage tracks were generated applying a mean windowing function with a smoothing window size of 2 pixels.

