## Supplementary material for "Transcription Represses Origin Activity in a Late-Replicating Fragile Site"

### Supplementary Fig.1.

(A) Crossing point (Cp) values obtained for $RT^+$-qPCR and $RT^-$-qPCR (background) experiments for relative quantification of nascent RNAs in the modified $DMD^{Tet/Tet}$ cell lines containing either the active or inactive minimal origin grown in the absence of tetracycline (as shown in Figure 1D) are reported in the table. NA indicates non-amplified signals. The amplicons used for quantification are shown in Figure 1B. (B) UCSC genome browser view showing the EU-seq signal across the $DMD^{Tet/Tet}$ gene carrying the active minimal origin on one allele in cells grown without and with tetracycline. The insertion site for the active minimal origin is indicated by a red dotted line. Below schematic representations of the inserted construct is shown. Black lines indicate amplicons used to distinguish the RT profile of the modified allele (With) and the WT allele (Without). (C) Replication timing (RT) profiles of each chromosomal allele were determined in the cell line described in A, using the allele-specific quantitative PCR analysis. The cells were grown in the absence of tetracycline before RT analysis. BrdU pulse-labeled cells were sorted into four S-phase fractions, from early to late (S1 to S4), and the immunoprecipitated newly synthesized DNA strands (NS) were quantified in each fraction by qPCR. Specific primer pairs were used to distinguish the RT profile of the modified allele (With), the WT allele (Without) and 5 kb away from the insertion site on both alleles (Both). The endogenous β-globin locus was analysed as an early-replicated control (Early).

### Supplementary Fig.2. PCR validation of clones selected for homologous recombination

(A-B) Schematic diagrams showing genomic regions containing a site-specific integrated construct. The 5′and/or 3′ arms of the targeted vector are shown as light grey boxes. Red arrows indicate primer pairs used to analyse the correct integration of the constructs by homologous recombination in the target region. PCR products were subjected to electrophoresis in a 1% w/v agarose gel and stained with SYBR safe. The DNA size marker is a commercial 1 kb DNA ladder (M). Lanes marked with a red star correspond to clonal cell lines selected for further analysis. (A) $DMD^{Tet/Tet}$ locus insertion site containing the minimal active or inactive origin is shown. (B) Late 2 insertion site, containing the minimal active or inactive origin associated with the β-actin+BlsR construct, is shown.

### Supplementary table 1 Primer sets used for plasmids constructions and quantitative PCR

Suppl Fig1

# A

| Crossing point values (Cp) | | *Bu1a* | *IL2R (2)* | *2xUSF (5')* | *Without* |
|---|---|---|---|---|---|
| RT$^+$-qPCR | Active minimal origin | 25,83 | 30,99 | 32,58 | 30,77 |
| | | 25,98 | 31,19 | 31,48 | 30,47 |
| | | 26,03 | 31,64 | 32,95 | 31,5 |
| | | 26,15 | 31,19 | 32,77 | 31,69 |
| RT$^-$-qPCR | Active minimal origin | 37,62 | 37,55 | NA | NA |
| | | NA | 40,21 | NA | NA |
| RT$^+$-qPCR | Inactive minimal origin | 26,85 | 31,85 | 33,61 | 31,21 |
| | | 26,48 | 31,59 | 33,28 | 31,46 |
| | | 27,22 | 32,09 | 33,68 | 31,84 |
| | | 26,46 | 32,49 | 33,97 | 31,43 |
| RT$^-$-qPCR | Inactive minimal origin | 45 | NA | NA | NA |
| | | NA | NA | NA | NA |

# B

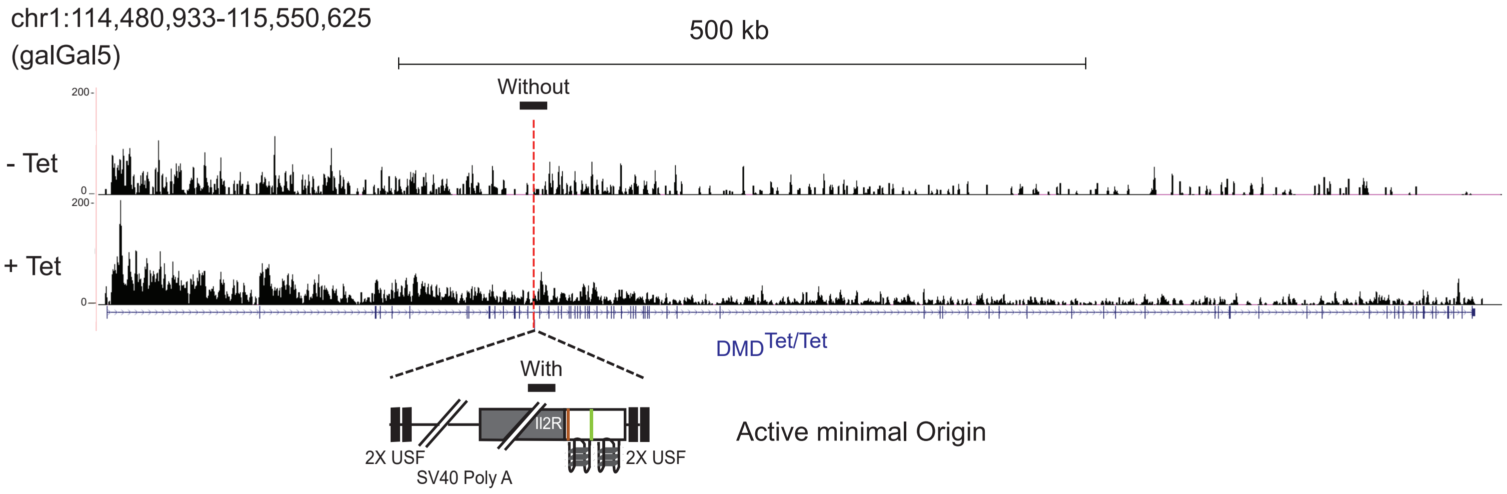


# C

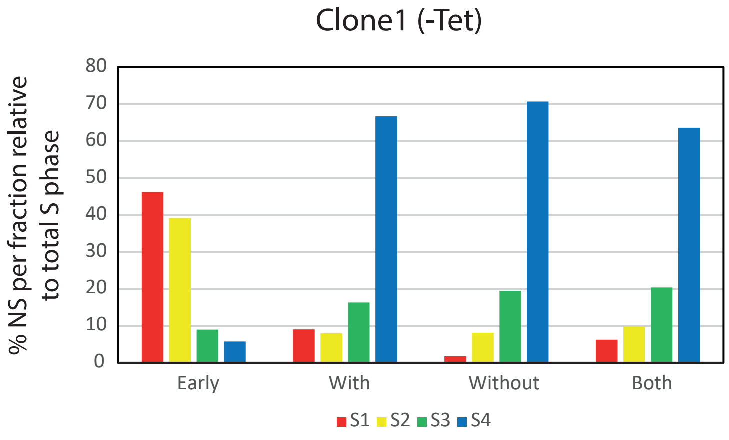

Suppl. Fig. 2

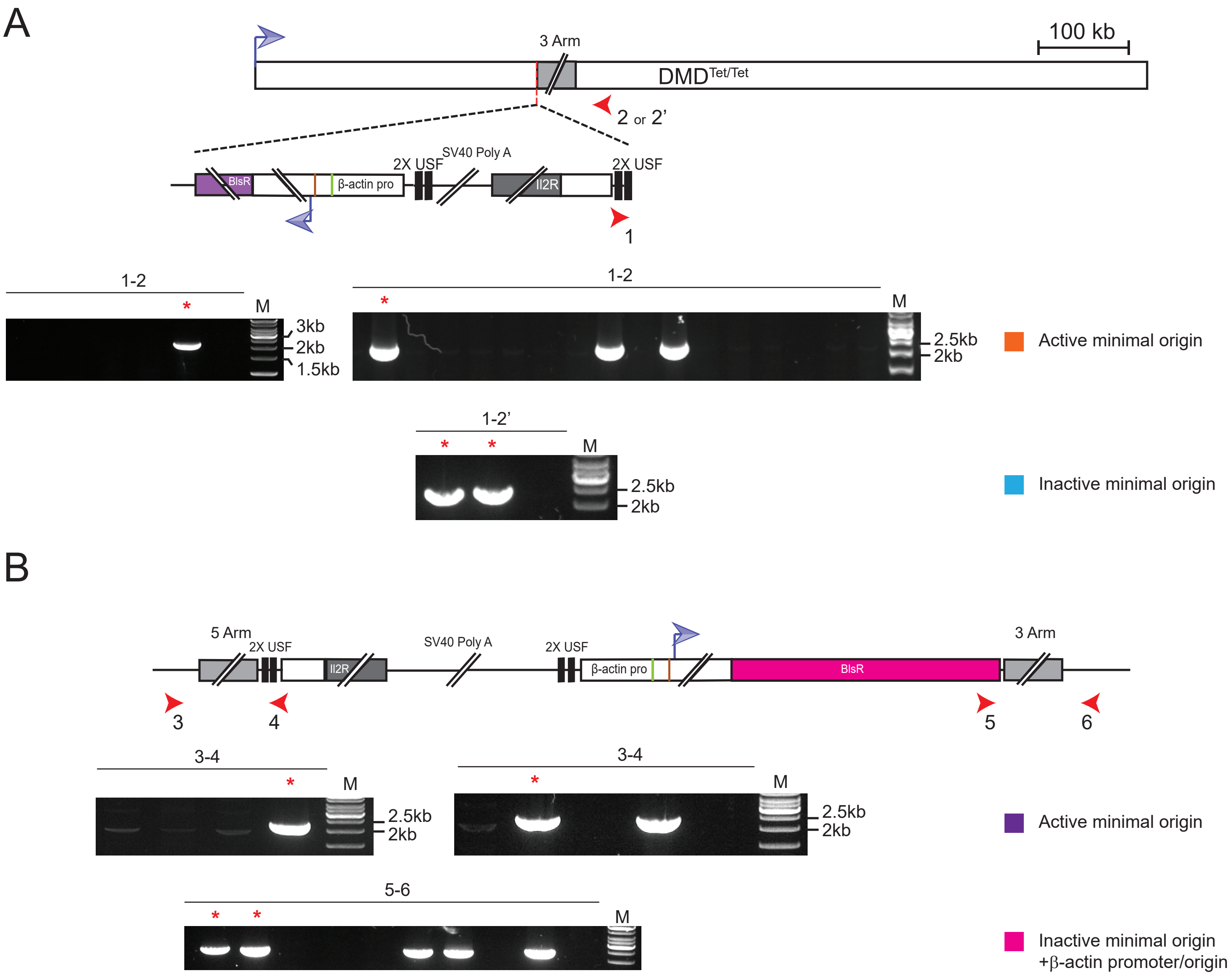
A
3 Arm
100 kb
DMD Tet/Tet
2 or 2'
BlsR
β-actin pro
2X USF
SV40 Poly A
Il2R
2X USF
1
1-2
M
3kb
2kb
1.5kb
1-2
M
2.5kb
2kb
Active minimal origin
1-2'
M
2.5kb
2kb
Inactive minimal origin
B
5 Arm
2X USF
Il2R
SV40 Poly A
2X USF
β-actin pro
BlsR
3 Arm
3
4
5
6
3-4
M
2.5kb
2kb
3-4
M
2.5kb
2kb
Active minimal origin
5-6
M
Inactive minimal origin
+β-actin promoter/origin

Table 1: Primer sets used for plasmids constructions and quantitative PCR

| Primer name | Figure | Forward primer sequence | Reverse primer sequence | Position (Build Mars 2018) |
|---|---|---|---|---|
| IL2R (1) /With | 1, 2, 3, SupplFig1 | GGGGACTGCTCACGTTCATCA | AATGTGGCGTGTGGGATCTC | |
| IL2R (2) | 1, 3 | CTACACAGAGGTCCTGCTG | TGCCTGAGGCTTCTCTTCAC | |
| 2xUSF (3') | 1, 3 | TGCATTCTAGTTGTGGTTTGTCC | ACCGTCGACCAACTTTGTATAGA | |
| **CFS locus** | | | | |
| 2xUSF (5') | 1 | TGTGTAGTGACTGTACCTATGCTT | CCGTGCCACCTCAACTTTTGTAT | |
| Bkgd (CFS site) / Both | 1, 2, SupplFig1 | TGGATTTCCCAGCAGCTCTT | GCTTCCCAACCCCAACTTTT | chr1:116871770-116871835 |
| Without | 1, 2, SupplFig1 | ACCATGCTAAGATGATGACTGTGA | TGTGAAGCTATGCAATGTTCCAGT | chr1:116877422+116877511 |
| **Late locus** | | | | |
| β-actin | 3 | TGCAGAAATCGGAGGAAGAAGA | GAATTGCCGCTCCCACATGA | |
| Bkgd (late site) | 3 | CGTCAGAGTGGTGGTGAGAA | TCTTGCCCAAACCAAAAACA | chr1:179238582+179238693 |
| Both | 3 | CAAGGTTTCCACCCCTAAAGA | TGATGGATGTGGGAAGAGAAA | chr1:179246338+179246419 |
| Wt allele / Without | 3 | TTTACACTACTCCCACCCCTCG | TTGACCATATGCCACCAACACC | chr1:179250225+179250324 |
| **Controls** | | | | |
| ρ-origin | 1, 3 | GACGGTCAGGTTTGCCAAAG | TCCTGAGGATACGTTTTTCAG | chr1:197287850-197288114 |
| Bu1a gene | 1 | AATGTCCCCAAAATGAGCTG | CCTCTTTTTCCACCCTCCTC | chr1:93337763+93337907 |
| Early timing control | 2, 3, Suppl.Fig1 | GACGGTCAGGTTTGCCAAAG | TCCTGAGGATACGTTTTTCAG | chr1:197287850-197288114 |
| Mitochondrial DNA | 2, 3 | CATCCCATGCATAACTCCTG | GTAGTCCAGGCTTCACTTGA | chrM:541+731 |
| **Constructions** | | | | |
| 5 Arm for minimal origin+β-actin-PuroR insertion in DMDTetO | | GGGGACAAGTTTGTACAAAAAAGCAGGCTTCCTGCCTCCAAGTCTACACA | GGGGACAACTTTTGTATACAAAGTTGCCTTACCTGCCCCAAACTGC | chr1:116875053+116877153 |
| 3 Arm for minimal origin+β-actin-PuroR insertion in DMDTetO | | GGGGACAACTTTGTATAATAAAGTTGTGGAGCAACTACCCTGTCCT | GGGGACCACTTTGTACAAGAAAGCTGGGTCCACCCAACTTGCACACTGT | chr1:116879085+116881038 |
| **screening of targetted integration** | | | | |
| 5'-screening-DMD gene for active minimal origin+β-actin-PuroR (1-2) | Suppl.Table1 | TACACATTCACAGAGGAAAAGA | GCCACCTCAACTTTTGTATAC | |
| 5'-screening-DMD gene for inactive minimal origin+β-actin-PuroR (1-2') | Suppl.Table1 | TGCCAAAACATATTTCTTCTCC | GCCACCTCAACTTTTGTATAC | |
| 5'-screening-late2 site for minimal active origin+β-actin-BlsR (3-4) | Suppl.Table1 | ACTTTGCACAAGCTAAGGAACC | AAAAGTTGAGGTGGCACGGG | |
| 3'-screening-late2 site for minimal inactive origin+β-actin-BlsR (5-6) | Suppl.Table1 | TGCATTCTAGTTGTGGTTTGTCC | ATCTCTGCCTTCAAACCTTCAG | |
